# *C9orf72* polyGA knock-in mice exhibit mild motor and proteomic changes consistent with ALS/FTD

**DOI:** 10.1101/2025.05.30.656994

**Authors:** Carmelo Milioto, Mireia Carcolé, Matteo Zanovello, Mhoriam Ahmed, Raja S. Nirujogi, Daniel Biggs, Eszter Katona, Idoia Glaria, Almudena Santos, Anny Devoy, Pietro Fratta, Dario R. Alessi, Ben Davies, Linda Greensmith, Elizabeth M.C. Fisher, Adrian M. Isaacs

**Author notes:** Correspondence to AMI or EMCF or CM.

## Abstract

A GGGGCC repeat expansion in *C9orf72* is the most common genetic cause of amyotrophic lateral sclerosis (ALS) and frontotemporal dementia (FTD). The repeat expansion is translated into five different dipeptide repeat proteins: polyGA, polyGP, polyGR, polyAP and polyPR. To investigate the effect of polyGA, which is the most abundant dipeptide repeat protein in patient brains, we used CRISPR/Cas9 to insert 400 codon-optimized polyGA repeats immediately downstream of the mouse *C9orf72* start codon. This generated (GA)400 knock-in mice driven by the endogenous mouse *C9orf72* promoter, coupled with heterozygous *C9orf72* reduction. (GA)400 mice develop subtle pathology including mild motor dysfunction characterized by impaired rotarod performance. Quantitative proteomics revealed polyGA expression caused protein alterations in the spinal cord, including changes in previously identified polyGA interactors. Our findings show that (GA)400 mice are a complementary *in vivo* model to better understand C9ALS/FTD pathology and determine the specific role of single DPRs in disease.

## Introduction

Amyotrophic lateral sclerosis (ALS) is the most frequent motor neuron disease in adults, characterised by progressive muscle weakness and atrophy leading to death typically from respiratory failure with a few years of diagnosis. Frontotemporal dementia (FTD) is the second most frequent young-onset dementia, characterised by behavioural, language and cognitive impairment. Despite these differences, ALS and FTD share clinical, neuropathological, and genetic features. A hexanucleotide GGGGCC (G_4_C_2_) repeat expansion in the first intron of the *C9orf72* gene is the most common genetic cause of ALS and FTD, collectively termed C9ALS/FTD [1]–[3]. Three mechanisms have been proposed to induce C9ALS/FTD pathology: (i) reduced transcription of *C9orf72*, (ii) the presence of sense and antisense repeat-containing RNA, (iii) expression of aberrant dipeptide repeat (DPR) proteins encoded in six frames by the hexanucleotide repeat [4].

The presence of the hexanucleotide repeats gives rise to repeat-associated non-ATG initiated (RAN) translation, a non-canonical protein translation mechanism that does not require an ATG start codon [5]. RAN translation occurs in every reading frame, encoding five different DPRs: polyGA, polyGR, polyGP, polyPA and polyPR, which all form neuronal cytoplasmic inclusions in C9ALS/FTD patient brains [6], [7]. Since the exact contribution of each DPR to disease pathogenesis is poorly understood, investigation of DPRs within the context of C9ALS/FTD is essential. We and others have demonstrated that DPR proteins are toxic *in vivo* and *in vitro* [4], [8]. A recent study of the interactome of DPRs identified interacting partner proteins involved in a variety of functions, including protein translation, signal transduction pathways, protein catabolic processes, amide metabolic processes and RNA-binding [9]. Although the arginine-containing polyGR and polyPR proteins (R-DPRs) are the most toxic DPRs based on studies in several systems [10]–[17], considerable evidence also shows that DPRs not containing arginine induce toxicity. In particular, polyGA, the most abundant DPR species in patients, forms highly insoluble aggregates [18]–[21].

Additionally, polyGA expression has been associated with TDP-43 phosphorylation and aggregation in cellular and mouse models, and this is a key feature in most human cases of ALS [22]–[24]. Recently, polyGA overexpression in primary rat cortical neurons has been shown to promote loss of synaptic proteins and cause autophagy defects [25]. Furthermore, analysis of cellular models and patient brain tissue revealed that polyGA aggregates sequester proteasome components, molecular chaperones and other proteins involved with protein folding/degradation pathways and nucleocytoplasmic transport, suggesting downstream defects in protein quality control and degradation [26]–[29].

Several mouse models have been generated to elucidate how the presence of hexanucleotide repeats leads to neurodegeneration [30], [31]. Early studies showed that *C9orf72* homozygous knock-out mouse models, differently from heterozygous animals, develop severe autoimmunity and lymphatic defects, suggesting C9orf72 is involved in immune cell function [32]–[39]. Mouse models of *C9orf72* hexanucleotide repeat expansions have also been generated though AAV-mediated delivery [40], [41], or bacterial artificial chromosome (BAC) integration [42]–[45]. Although most of these mouse models recapitulate aspects of C9ALS/FTD pathology, there is considerable variation between them, including the chromosomal site expressing the repeats, suggesting more refined models are needed [46]. Additionally, several mouse models have been developed to better understand the role of specific DPRs. Studies of these mice show that *in vivo* expression of arginine-rich DPRs (R-DPRs) can drive toxicity [13]–[17], [47]. Similarly, studies in mice that over-express codon-optimised polyGA show that this can drive toxicity leading to neurodegeneration, motor and behavioural abnormalities, and inflammation [29], [48]. Furthermore, expression of polyGA induces selective neuron loss, interferon responses, and phosphoTDP-43 inclusions similar to those observed in C9ALS patients [49]. Finally, in both cell and mouse models, antibodies targeting polyGA could reduce polyGA, polyGP, and polyGR inclusions, improve behavioural deficits, decrease neuroinflammation and neurodegeneration, and increase survival [50]–[53].

To date, multiple mechanisms have been implicated as common downstream molecular pathways in C9ALS/ FTD, including autophagy, nucleocytoplasmic transport, pre-messenger RNA splicing, stress granule dynamics, DNA damage repair, mitochondrial dysfunction, nuclear pore alterations, impaired translation, lipid dyshomeostasis and synaptic dysfunction [4], [54]–[58]. However, despite considerable effort, the most relevant mechanism(s) by which the repeat expansion causes C9ALS/FTD remain unclear and no effective therapies have been developed to date. Thus, to understand the role(s) of individual DPRs *in vivo* and determine mechanisms associated with disease progression, we generated a series of *C9orf72* DPR knock-in mice. We recently described polyGR and polyPR knock-in mouse models that recapitulate many aspects of C9ALS/FTD pathology, such as cortical hyperexcitability and spinal motor neuron loss, and revealed a conserved neuroprotective extracellular matrix (ECM) signature in *C9orf72* ALS/FTD neurons [59]. Here we describe polyGA knock-in mice, in which 400 polyGA repeats were inserted directly after the mouse *C9orf72* ATG start codon, resulting in a mouse model that expresses polyGA repeats from the *C9orf72* locus combined with heterozygous *C9orf72* reduction. We carried out phenotypic analyses of this mouse on a C57BL/6J background (the same background as our polyGR and polyPR knock-in animals) and found a mild phenotype, suggesting polyGA expression at a physiological level leads to subtle changes that are consistent with, but less pronounced, than seen in other *in vivo* and *in vitro* C9ALS/FTD experimental models.

## Results

### Generation of *C9orf72* polyGA knock-in mice

In order to generate polyGA knock-in mice we adopted the same approach we used recently to generate polyGR and polyPR knock-in mice, which we will collectively refer to as R-DPR mice [59]. We first built patient-length, seamless, codon-optimised polyGA repeats by using recursive directional ligation [10], [60] and flanked them with epitope tags (**Fig. 1A**). We then used CRISPR-Cas9 technology to insert the repeats directly after the endogenous mouse *C9orf72* ATG start codon in mouse embryonic stem (ES) cells (**Supplementary Fig. 1A**).

**Figure 1.**
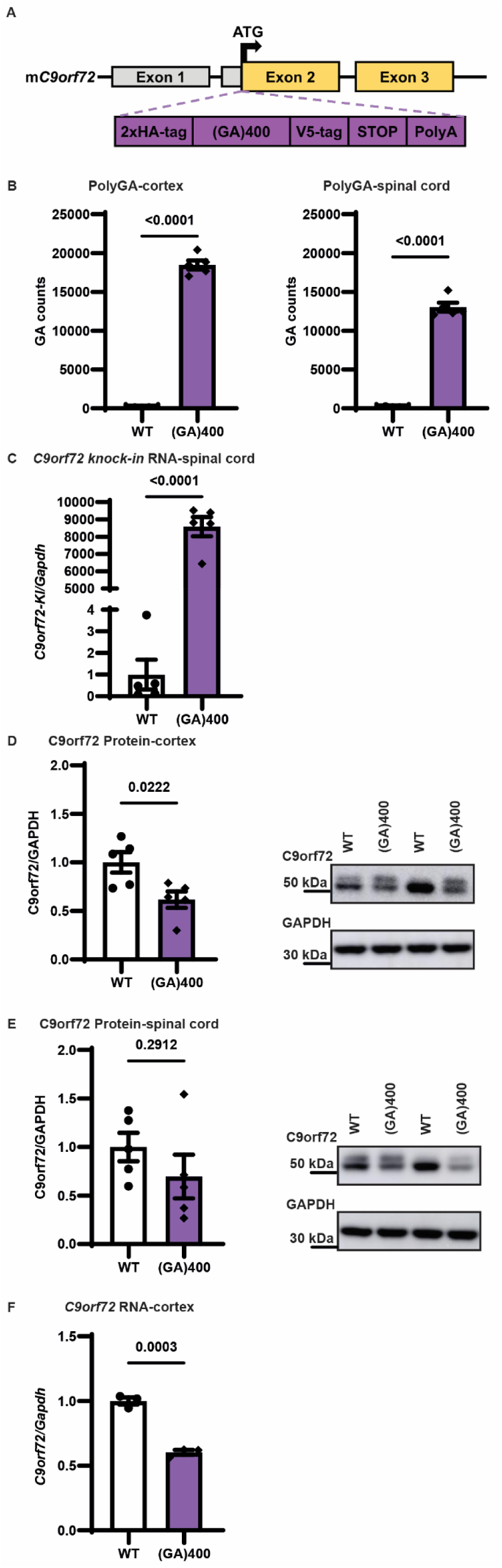
Generation of *C9orf72* polyGA knock-in mice. (A) Targeting strategy to generate (GA)400 mice with the knock-in sequence inserted in exon 2 of mouse *C9orf72* immediately after, and in frame with, the endogenous ATG. Schematic shows the genomic region and the knock-in targeting construct. Exons are shown boxed, untranslated regions of exons are coloured grey with translated regions in yellow. The targeting construct (in purple) contains the knock-in sequence composed of a double HA-tag, 400 codon-optimised GA repeats, a V5-tag, a stop codon, and a 120 bp polyA tail. (B) Quantification of polyGA proteins in cortex (left panel) and spinal cord (right panel) of WT and (GA)400 mice at 12 months of age by Meso Scale Discovery (MSD) immunoassay. Graph, mean ± SEM, n = 5 mice per genotype, two-sided unpaired two-sample Student’s t-test. (C) qPCR analysis of *C9orf72* knock-in transgene transcript levels normalised to *Gapdh* in spinal cord of WT and (GA)400 mice at 12 months of age. Common primers in the HA-tag and upstream endogenous mouse sequence were used to enable direct comparison between all lines. Graph, mean ± SEM, n = 5 mice per genotype, two-sided unpaired two-sample Student’s t-test. (D) Western blotting analysis of C9orf72 protein levels in cortex of WT and (GA)400 mice at 12 months of age. GAPDH is shown as loading control. Graph, mean ± SEM, n = 5 mice per genotype, two-sided unpaired two-sample Student’s t-test. (E) Western blotting analysis of C9orf72 protein levels in spinal cord of WT and (GA)400 mice at 12 months of age. GAPDH is shown as loading control. Graph, mean ± SEM, n = 5 mice per genotype, two-sided unpaired two-sample Student’s t-test. (F) qPCR analysis of *C9orf72* transcript levels normalized to *Gapdh* in brain cortex of WT and (GA)400 at 9 months of age. Graph, mean ± s.e.m.; n = 3 mice per genotype; two-sided unpaired two-sample Student’s t-test.

We next used targeted locus amplification to identify targeted ES cells that had a single insertion site, correct targeting, and no backbone co-integration (**Supplementary Fig. 1B-1D**). Validated ES cell clones were then used to generate knock-in mice using standard procedures. Briefly, targeted ES cells (C57BL/6N) were injected into C57BL/6J blastocysts, and the resulting chimeras were crossed to wild-type C57BL/6J mice for germline transmission, founders were backcrossed to C57BL/6J animals for a minimum of 5 generations before phenotypic analysis.

Heterozygous knock-in mice and littermate controls were aged and assessed, initially for polyGA expression. We detected polyGA in brain and spinal cord of (GA)400 animals at 12 months of age using an MSD immunoassay (**Fig. 1B**). As expected, at the same age, (GA)400 mice selectively expressed the knock-in sequence mRNA (**Fig. 1C**). As shown previously with our R-DPR knock-in mice [59], (GA)400 mice exhibited a significant ∼40% reduction in C9orf72 at both mRNA and protein levels in cortex and spinal cord compared to wildtype littermates (WT) mice (**Fig. 1D-F**). These results show that our (GA)400 mouse line expresses polyGA, alongside *C9orf72* reduction.

### (GA)400 mice develop mild motor dysfunction

We next evaluated the motor function of (GA)400 mice. Monthly body weight measurements indicated that (GA)400 male and female animals did not differ from WT littermates up to 18 months of age (**Fig. 2A**). Next, we assessed motor coordination by conducting rotarod analysis from 3 to 18 months of age. Male (GA)400 mice performed significantly worse than their WT littermates in accelerated rotarod performance from 6 months of age, with female mice showing a similar but non-significant trend in the same direction (**Fig. 2B**). Additionally, we performed grip strength analysis and neither (GA)400 male nor female knock-in mice showed significant strength deficits up to 18 months of age, although (GA)400 males consistently performed less well than their littermate controls (**Fig. 2C**). Based on these results, we suggest that (GA)400 mice develop a milder motor dysfunction as they age, as compared to our previously characterised R-DPR knock-in mouse models [59].

**Figure 2.**
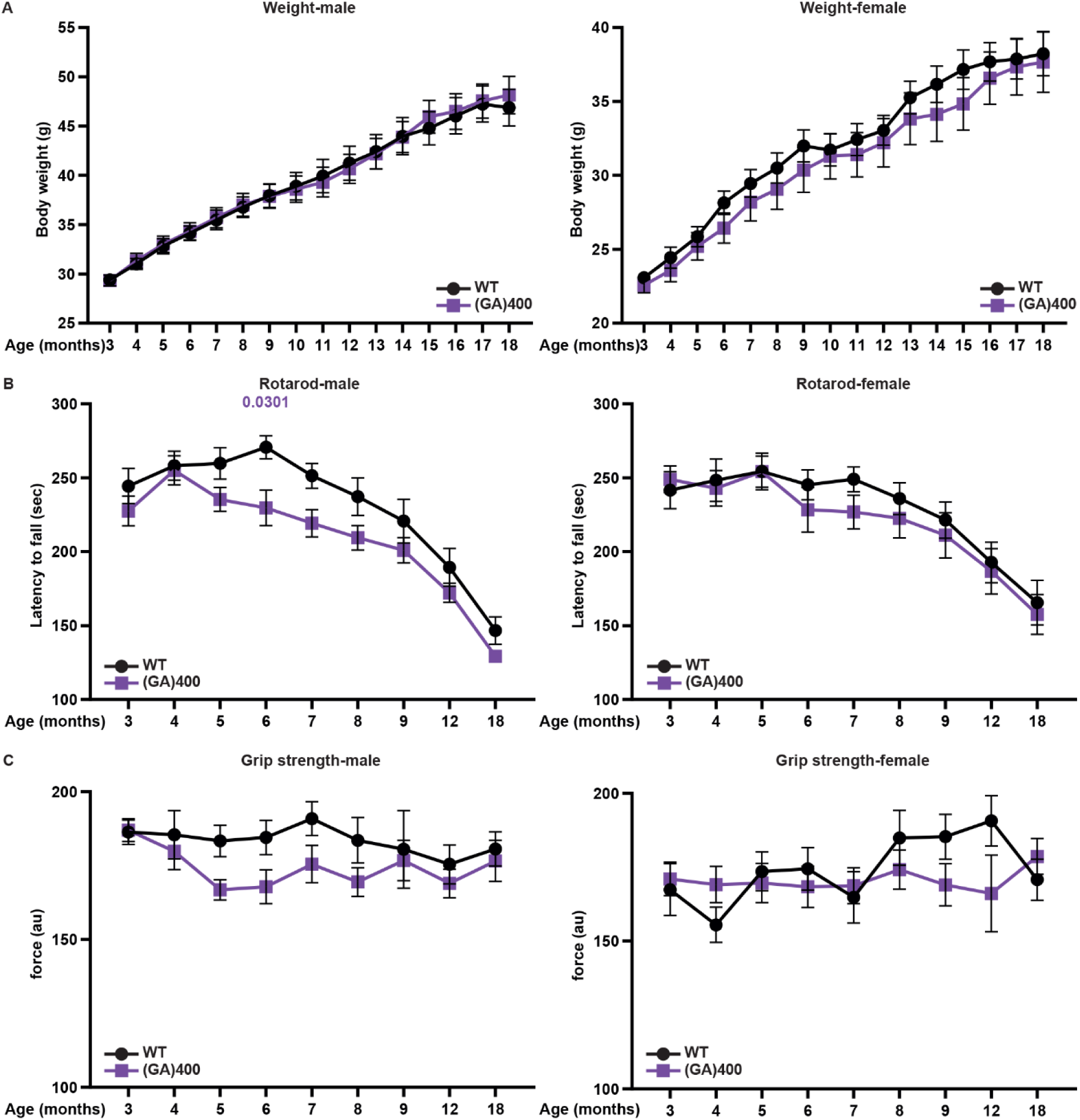
(GA)400 mice develop mild motor dysfunction. (A) Body weights of WT and (GA)400 male (left panel) and female (right panel) mice up to 18 months of age. Graph, mean ± SEM, n = 14 mice per genotype, two-way ANOVA, Bonferroni’s multiple comparison. (B) Accelerated rotarod analysis of motor coordination in WT and (GA)400 male (left panel) and female (right panel) mice up to 18 months of age. Graph, mean ± SEM, n = 14 mice per genotype, two-way ANOVA, Bonferroni’s multiple comparison. (C) Grip strength analysis of muscle force in WT and (GA)400 male (left panel) and female (right panel) mice up to 18 months of age. Graph, mean ± SEM, n = 14 mice per genotype, two-way ANOVA, Bonferroni’s multiple comparison.

### PolyGA expression does not cause overt pathology up to 18 months of age

We next investigated whether the subtle, but prolonged motor phenotype could be associated with motor neuron loss. We performed motor neuron counts at 9, 12, and 18 months of age in the ventral horn of the lumbar spinal cord as this is where the large alpha motor neurons reside that are primarily affected in ALS. Notably, we did not observe significant alteration in motor neuron numbers, but there was a non-significant trend towards lower motor neuron numbers as the mice aged compared to WT mice (**Fig. 3A-D**). In our R-DPR knock-in mice we observed motor neuron loss at 12 months of age that was confirmed using electrophysiological motor unit number estimation (MUNE) analysis [59]. To provide a direct comparison, we performed MUNE analysis in the hindlimbs of 12-month-old (GA)400 mice. Unlike our R-DPR knock-in mice, we did not observe a reduction in functional motor unit number in the extensor digitorum longus (EDL) muscles of (GA)400 mice when compared with WT littermates (**Supplementary Fig. 2A**). Additionally, we investigated whether polyGA expression causes other ALS/FTD pathological hallmarks. We conducted immunostaining analysis in 18-month-old animals and did not observe microgliosis in the lumbar spinal cord of (GA)400 mice (**Fig. 3E**). Similarly, we did not detect signs of astrogliosis in the lumbar spinal cord of 18-month-old (GA)400 mice (**Fig. 3F**).

**Figure 3.**
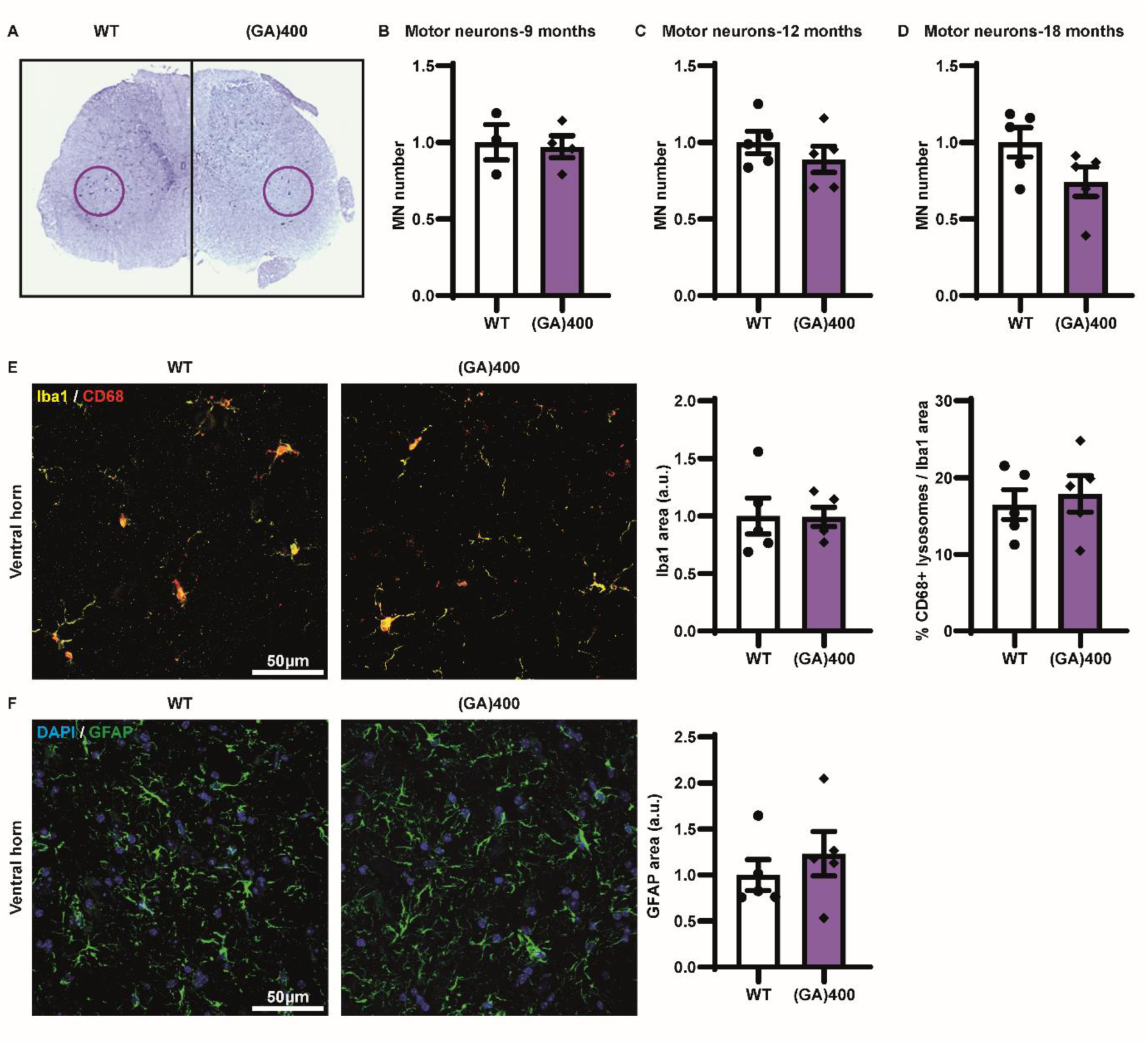
PolyGA expression causes modest age-dependent spinal motor neuron loss. (A) Panel shows representative image of Nissl staining of lumbar spinal cord in WT and (GA)400 mice at 18 months of age; the purple line delineates the sciatic motor pool in which motor neurons were counted. (B) Quantification of Nissl-stained motor neurons (MN) in lumbar spinal cord region L3-L5 in WT and (GA)400 mice at 9 months of age. Graph, mean ± SEM, n = 3-4 mice per genotype, two-sided unpaired two-sample Student’s t-test. (C) Quantification of Nissl-stained motor neurons in lumbar spinal cord region L3-L5 in WT and (GA)400 mice at 12 months of age. Graph, mean ± SEM, n = 5 mice per genotype, two-sided unpaired two-sample Student’s t-test. (D) Quantification of Nissl-stained motor neurons in lumbar spinal cord region L3-L5 in WT and (GA)400 mice at 18 months of age. Graph, mean ± SEM, n = 5 mice per genotype, two-sided unpaired two-sample Student’s t-test. (E) Representative confocal images and quantification of immunofluorescence staining showing microglial density and colocalization between microglial markers Iba1 (yellow) and microglial lysosomal marker CD68 (red) in lumbar spinal cord ventral horn in WT and (GA)400 mice at 18 months of age. Graph, mean ± SEM, n = 5 mice per genotype, two-sided unpaired two-sample Student’s t-test. (F) Representative confocal images and quantification of immunofluorescence staining of astrocytic marker GFAP (green) in lumbar spinal cord ventral horn in WT and (GA)400 mice at 18 months of age. DAPI (blue) stains nuclei. Graph, mean ± SEM, n = 5 mice per genotype, two-sided unpaired two-sample Student’s t-test.

### PolyGA expression induces proteomic alterations in the spinal cord

We performed quantitative proteomics in the lumbar spinal cord of 12-month-old (GA)400 mice, which revealed modest changes at the protein level (**Table 1**). Among these alterations were reduced levels of C9orf72 and its binding partner SMCR8 (**Fig. 4A**), which confirms the reduced C9orf72 levels we observed by western blotting (**Fig 1E**).

**Figure 4.**
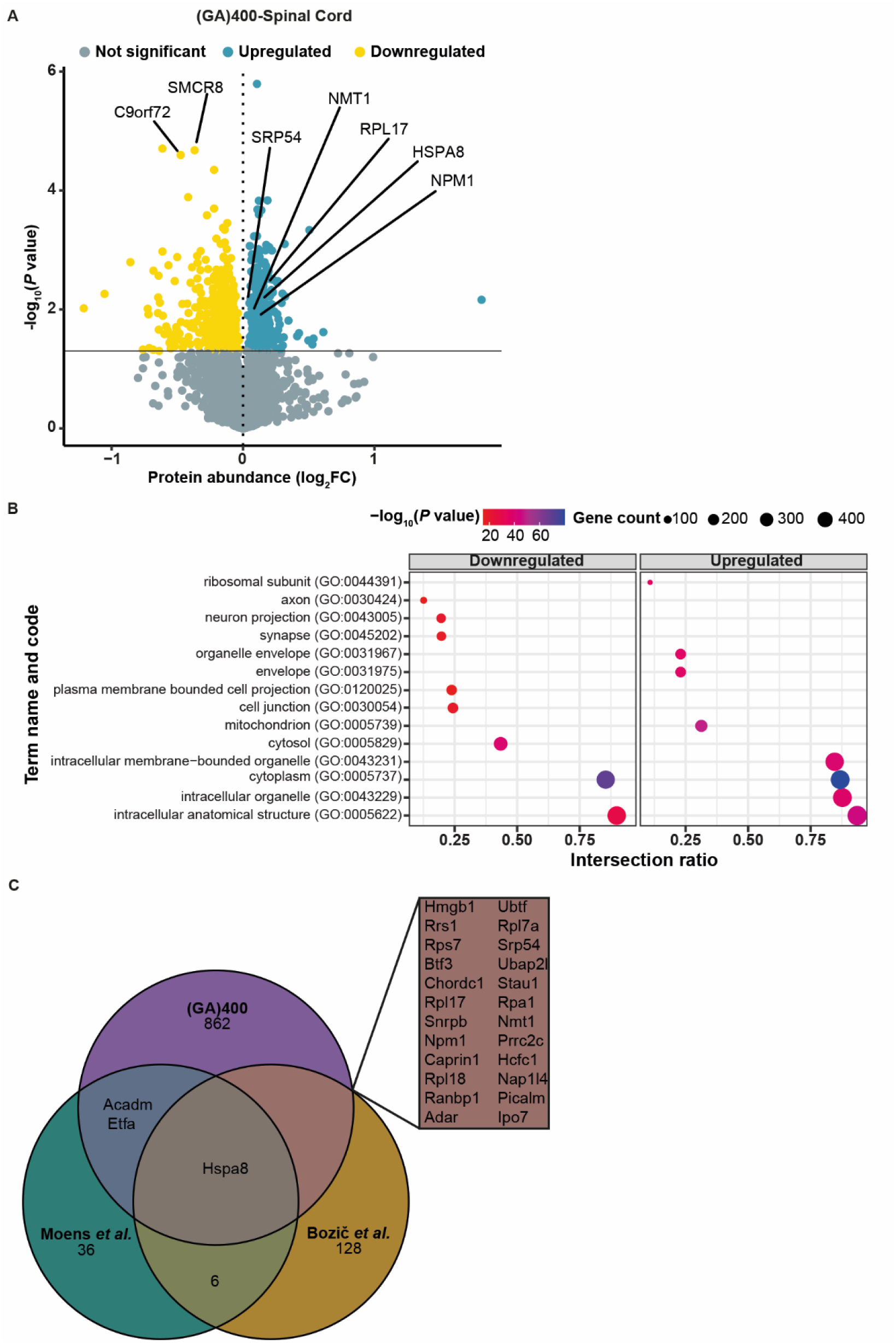
PolyGA expression induces proteomic changes in the spinal cord. (A) Protein expression volcano plots from the lumbar spinal cord of 12-month-old (GA)400. n = 5 mice per genotype, two-sided Welch’s t-Test with 5% FDR multiple-correction. (B) Significantly enriched downregulated (left) and upregulated (right) Gene Ontology (GO) pathways in the lumbar spinal cord of 12-month-old (GA)400 mice. Proteomics performed on n = 5 mice per genotype, two-sided Welch’s t-Test with 5% FDR multiple-correction. The x-axis represents the intersection ratio between significant genes and the total number of genes for the specific GO category, while the size and colour of dots represent the total count and-log10(p-value) of significant genes for each category, respectively. (C) Venn diagram showing the overlap of GA(400) significant targets with the orthologues of the significant hits from *Moens et al., 2019* [58] and *Bozič et al., 2022* [9].

We previously observed an upregulation of ECM components in R-DPR knock-in mouse lumbar spinal cord at 12 months of age using the same quantitative proteomics approach [59]. However, ECM proteins were not significantly increased in (GA)400 mice at 12 months of age (**Supplementary Fig. 3A**). As COL6A1 was the most increased ECM component in our R-DPR knock-in mice, we measured COL6A1 levels in the (GA)400 mice using immunoblotting. We did not detect alterations in COL6A1 protein level in cortex or spinal cord of 12-month-old (GA)400 mice (**Supplementary Fig. 3B**). Similarly, COL6A1 volume in spinal cord of 12-month-old (GA)400 mice was not altered (**Supplementary Fig. 3C**).

Gene Ontology (GO) molecular processes term enrichment analysis revealed several significantly downregulated pathways in lumbar spinal cord of (GA)400 mice, including synapse proteins (**Fig. 4B**), which were similarly downregulated in R-DPR knock-in spinal cord. Additionally, we compared our proteomic changes to recently published polyGA interactome studies in cells [9] and fruit flies [58] (**Fig. 4C**). We found that HSPA8, a molecular chaperone previously associated with poly-GA aggregates and TDP-43 toxicity [26], [61], was influenced by polyGA expression in all three models analysed. Notably, additional polyGA interactors, including NPM1, NMT1, RPL17, and SRP54 were significantly upregulated within our (GA)400 proteomic dataset (**Fig. 4C**). This analysis suggests that physiological polyGA expression affects known polyGA interactor proteins *in vivo*. Thus, our polyGA knock-in mice exhibit changes observed in other models and may provide insight into early pathological changes caused by polyGA.

## Discussion

Here, we describe a novel knock-in mouse to further investigate the role of DPR toxicity in C9ALS/FTD pathology. Using the same approach we previously adopted for R-DPR mice [59], we generated a mouse model that selectively expresses polyGA from the endogenous locus, with a reduced level of C9orf72, thus recapitulating two key features of *C9orf72* ALS/FTD. Briefly, we inserted a codon optimised polyGA sequence immediately after and in frame with the mouse *C9orf72* ATG start codon. This strategy permits us to take advantage of the endogenous mouse *C9orf72* promoter and also removes one *C9orf72* allele, thus reducing C9orf72 levels. Notably, we intentionally generated mouse models that focus on DPRs in isolation. This means they do not model RAN translation or RNA toxicity, which will require other approaches. On the other hand, our reductionist approach allows a direct comparison of the effects of polyGA, polyGR, and polyPR repeats in *in vivo* mouse models on a common genetic background.

(GA)400 mice exhibited a deficit in rotarod performance that was less prominent than in our (GR)400 and (PR)400 mice. Similarly, (GA)400 mice showed a non-significant trend towards motor neuron loss at 12 and 18 months of age, while (GR)400 and (PR)400 mice showed significant reductions in motor neurons at these timepoints. We interpret these results as showing that polyGA is not benign but is likely better tolerated and thus leads to milder effects than the R-DPRs, at least in the first 18 months of life in our mouse model. This supports evidence from several experimental models where the arginine-rich polyGR and polyPR have been shown to exert stronger neurotoxic effects than polyGA [3], [10], [30].

Based on our recent findings that expression of R-DPRs induces increased levels of specific ECM proteins in the spinal cord, which provide protection against neurodegeneration [59], we focused on spinal cord to identify potential mechanisms associated with polyGA expression. We performed proteomics analysis in the spinal cord at 12 months of age and analysed whether (GA)400 mice show the ECM signature we described in R-DPR mice and *C9orf72* iPSC-derived motor neurons. However, no significant change in ECM proteins was observed. Once more, these results are consistent with the milder effects observed in polyGA knock-in mice.

While the increased ECM signature was the most striking proteomic change in (GR)400 and (PR)400 mice, we also reported downregulated pathways with lower fold-changes. Due to the modest toxicity of polyGA, we did not expect proteomics analysis to reveal a strong similarity between (GA)400 and R-DPR mice. Surprisingly, we found several GO terms related to downregulated genes in the (GA)400 mouse that were shared with the R-DPR knock-in mouse models. Notably, these GO terms are related to neuronal and cellular homeostasis including synapse, neuron projection, plasma membrane bound cell projection, cell junction, and axon, indicating neuronal/axonal or synaptic dysfunction. This observation suggests that each DPR causes neuronal dysfunction, even if GR and PR appear to be more toxic over the 18-month time frame of our studies.

Consistent with other knock-in models, the phenotypes were relatively modest across all our DPR knock-in mouse lines. This suggests that other factors are necessary in addition to DPR expression to drive the overt neuronal loss observed in patients. One possibility is an age-related decline in proteostasis that may allow higher levels of DPRs to accumulate over time, potentially closer to the levels utilised in over-expression studies that show overt toxicity and/or TDP-43 mislocalisation [24], [47], [62].

Since we and others have previously performed polyGA interactome studies [9], [58], we compared results obtained from these studies with our proteomics data. This comparison revealed that NPM1, NMT1, RPL17, SRP54, and HSPA8 were common across the datasets analysed. Interestingly, each of these proteins were upregulated in (GA)400 spinal cord. This could suggest a stabilising interaction with polyGA, but further studies will be required to confirm this and whether there are functional consequences. Notably, this conservation of events across several models indicates a genuine phenomenon linked to the presence of polyGA.

Overall, our study shows that our novel (GA)400 mouse model exhibits mild motor and proteomic changes that may model the earliest effects of polyGA in the brain and spinal cord. Together with our (GR)400 and (PR)400 mice, this series of DPR *C9orf72* knock-in mice provides a valuable tool to study the role of DPRs i*n vivo*.

## Acknowledgements

We thank the staff of the MRC Prion Unit at UCL Biological Services Facility; Nick Kaye, Craig Fitzhugh, Juliette Ajok-Omona, and Rebekah Hallpike for experimental assistance. We also thank the UK DRI at UCL technicians Elena Ghirardello and Phill Muckett, and the Biological Services Units at UCL.

This work was supported by the Motor Neuron Disease Association and the UK Dementia Research Institute through UK DRI Ltd, principally funded by the Medical Research Council

## Author Contributions Statement

Design of the study and conceptualization: C.M., E.M.C.F., and A.M.I. Experiments and data generation: C.M., M.C., M.Z, M.A., R.S.N, D.B., E.K., I.G., A.S. Supervised research: A.D., P.F., D.R.A., B.D., L.G. Writing of the original and revised draft: C.M., E.M.C.F., and A.M.I. All authors read and approved the manuscript.

---

**Supplementary Figure 1.**
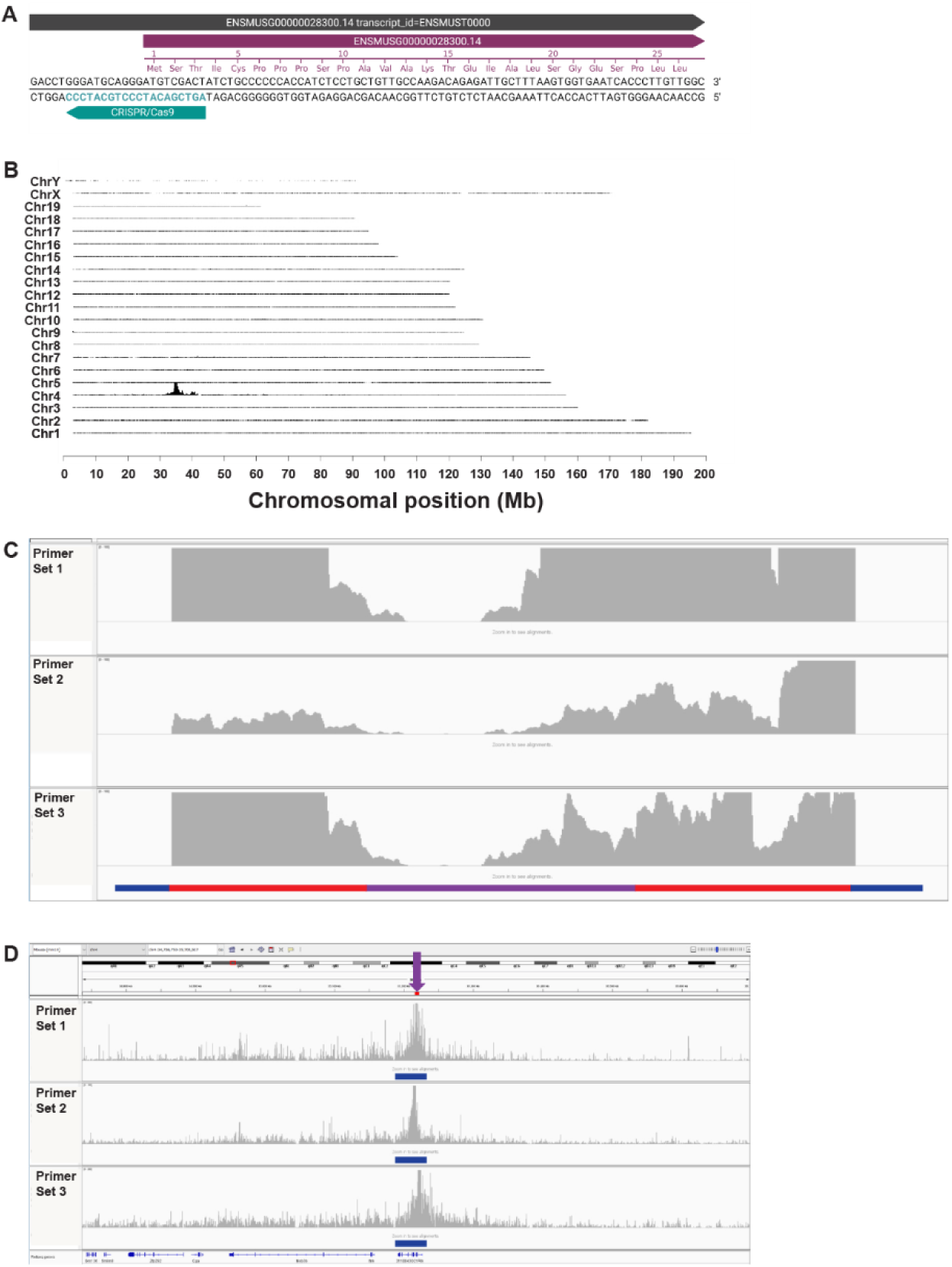
*C9orf72* knock-in strategy and confirmation. (A) Design for CRISPR assisted *C9orf72* gene targeting. The sgRNA for CRISPR/Cas9 is indicated by the teal bar. (B) Mapping of targeted locus amplification reads across the mouse genome. The chromosomes are indicated on the y-axis, the chromosomal position on the x-axis. (C) Targeted locus amplification sequence coverage across the knock-in sequence. The whole knock-in sequence has good coverage except for the blue underlined backbone sequences as expected from a correct targeting event as the backbone is not included. A coverage gap is present at the location of the 400 polyGA repeats, and is underlined by the purple bar. The red bars mark the homology arms. Y-axis is limited to 100x.Y-axis is limited to 100x. (D) Targeted locus amplification sequence coverage across the knock-in integration locus. The red bar marks the homology arms. The blue bar marks the Refseq Genes (3110043O21Rik). The purple arrow indicates the knock-in integration site. Y-axes are limited to 500x, 100x and 200x respectively.

**Supplementary Figure 2.**
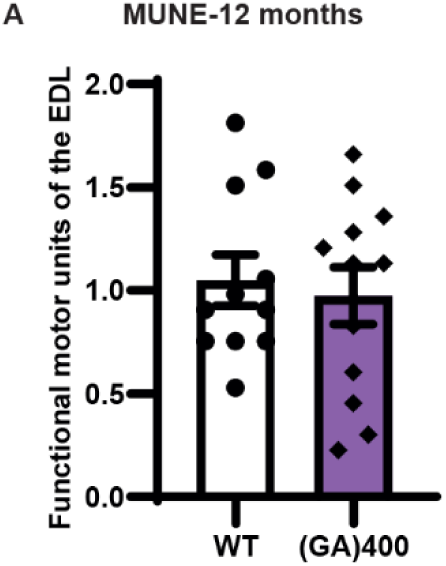
(GA)400 knock-in mice do not exhibit functional motor unit alteration at 12 months of age. (A) Quantification of MUNE determined in EDL muscle in WT and (GA)400 mice at 12 months of age. Graph, mean ± SEM, n mice = 6 WT and 6 (GA)400, n muscles = 11 WT and 12 (GA)400, two-sided unpaired two-sample Student’s t-test.

**Supplementary Figure 3.**
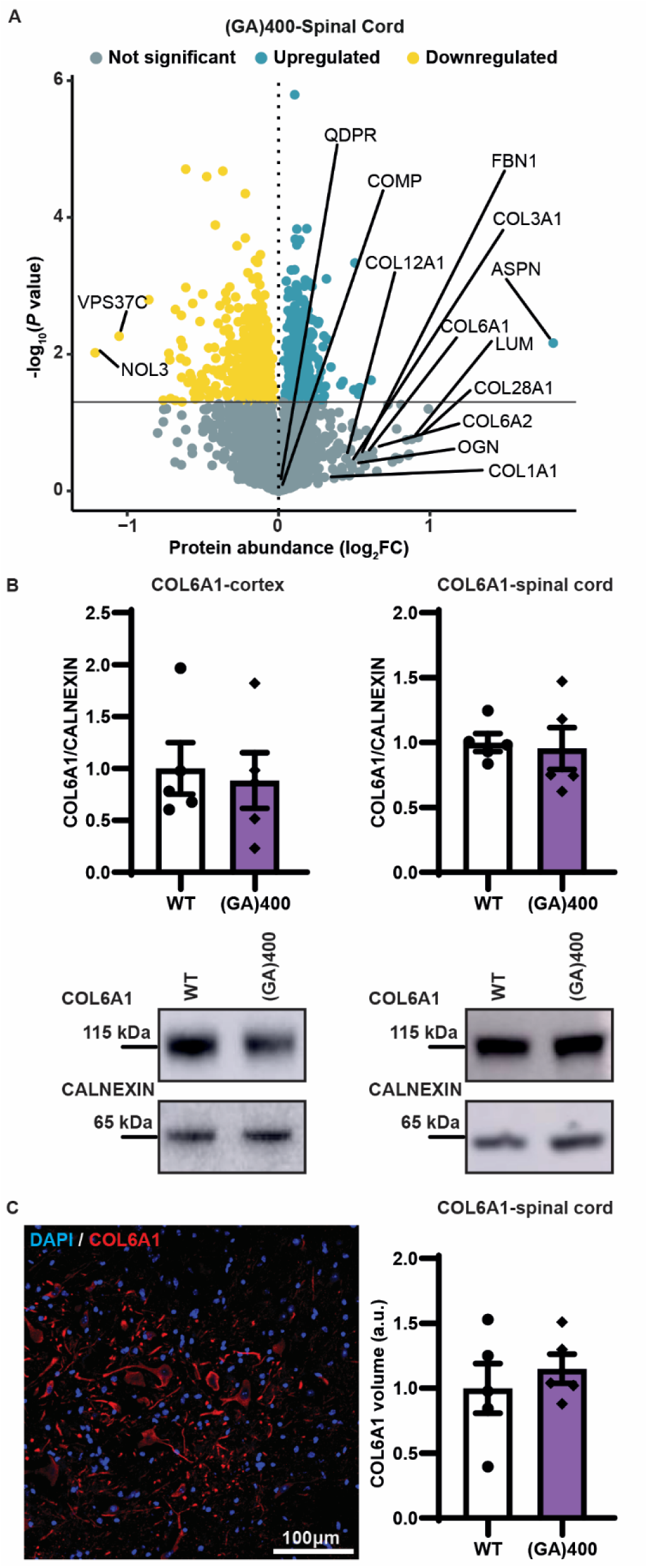
(GA)400 knock-in mouse spinal cord does not show increased extracellular matrix protein levels. (A) Protein expression volcano plots from the lumbar spinal cord of 12-month-old (GA)400. N = 5 mice per genotype, two-sided Welch’s t-Test with 5% FDR multiple-correction. (B) Western blot of COL6A1 in cortex (left panel) and lumbar spinal cord (right panel) of WT and (GA)400 mice at 12 months of age. Calnexin is shown as loading control. Graph, mean ± SEM, n = 5 mice per genotype, two-sided unpaired two-sample Student’s t-test. (C) Representative confocal image and quantification of immunofluorescence staining of astrocytic marker COL6A1 (red) in lumbar spinal cord ventral horn in WT and (GA)400 mice at 12 months of age. DAPI (blue) stains nuclei. Graph, mean ± SEM, n = 5 mice per genotype, two-sided unpaired two-sample Student’s t-test.

## Methods

### Assembly of targeting constructs

Assembly of targeting constructs was performed as previously described [59]. Briefly, 100 codon-optimised polyGA repeats were synthesized (Thermo Fisher Scientific). A pMC cloning vector was synthesized to contain 600 bp of the 5’ homology arm, a double HA-tag, a V5 epitope tag, a SV40 polyA tail, and 250 bp of 3’ homology arm. 100 codon-optimised DPRs were cloned into this vector within the *BbsI* and *BsmBI* sites to generate pMC-(GA)100. 400 codon-optimised DPRs were assembled with two consecutive rounds of recursive directional ligation taking advantage of the restriction enzymes *BbsI* and *BsmBI* to generate pMC-(GA)400.

A selection cassette (FRT-PGK-gb2-neo-FRT, Gene Bridges) was inserted in the pMC-(GA)400 in the *NheI* site. A BAC subcloning kit (Red/ET recombination, Gene Bridges) was used to clone full length homology arms (2.7 kb in 5’ and 3.2 kb in 3’) from the BAC clone (RP23-434N2), containing the C57BL/6J sequence of the mouse *C9orf72* gene, into the targeting vector pBlueScript II SK (+). Knock-in constructs were obtained by inserting sequences from pMC-(GA)400 into targeting vectors within the *BstXI* and *XcmI* sites.

### Animals

All procedures involving mice were conducted in accordance with the Animal (Scientific procedures) Act 1986, were performed at University College London under an approved UK Home Office project licence reviewed by the Institute of Prion Diseases Animal Welfare and Ethical Review Body and were reported according to the ARRIVE2 guidelines. Mice were maintained in a 12 h light/dark cycle at a temperature of 20°C-24°C and relative humidity of 45%-55% with food and water supplied *ad libitum*. The knock-in mice generated are available from the European Mutant Mouse Archive, EMMA ID EM:14658, strain name B6J.B6N-C9orf72^em1.1(GR400)Aisa^/H.

Assembly of targeting constructs was performed as previously described [59]; to generate the (GA)400 mouse strain, we performed CRISPR assisted gene targeting in JM8F6 embryonic stem (ES) cells (C57BL/6N) using our targeting vector(s) and a CRISPR/Cas9 guide designed against the insertion site.

The CRISPR construct (pX330-Puro-C9orf72), expressing Cas9 and a U6 promoter driven single-guide-RNA (sgRNA) designed against the following sequence AGTCGACATCCCTGCATCCC. This was generated by annealing the following two oligos (5’-CACCgAGTCGACATCCCTGCATCCC-3’; 5’-AAACGGGATGCAGGGATGTCGACTc-3’) and cloning this into the unique *BbsI* sites of pX330-U6-Chimeric_BB-CBh-hSpCas9 (Addgene # 42230), modified by the addition of a PGK-Puro cassette.

1×10^6^ ES cells were electroporated with 2.5 µg of the cloned pX330-Puro-C9orf72 plasmid and 2.5 µg of targeting vector using the Neon Transfection System (Thermo Fisher Scientific) (3 × 1400 V, 10 ms) and plated on puromycin resistant fibroblast feeder layers. After approximately 24 hours, 600 ng/ml puromycin was applied for a further 48 hour to allow transient selection. After further 5 days in culture without selection, individual colonies were isolated, expanded and screened for the desired targeting at both the 5’ end (5’-TCGGGGATTATGCCTGCTGC’3’ and 5’-GCATCCCAGGTCTCACTGCA-3’) and at the 3’ end (5’-TCGAAAGGCCCGGAGATGAGGAAG-3’ and 5’-GGGTTCAGACAGGTACAGCAT-3’). ES cells from correctly targeted clones were injected into albino C57BL/6J blastocysts and the resulting chimeras were mated with albino C57BL/6J females. The presence of the targeted allele in the F1 generation was confirmed at the DNA level by the above PCR and Sanger sequencing. Germline-transmitting founders were obtained and backcrossed to wild-type C57BL/6J mice to maintain hemizygous lines for a minimum of 5 generations.

Mouse genotype was determined by PCR for knock-in sequence with the following set of primers (forward 5’-TAAGCACAGCAGTCATTGGA-3’ and reverse 5’-AAGCGTAATCTGGAACATCG-3’). Repeat length was determined by PCR with the following set of primers (forward 5’-CCCATACGATGTTCCAGATTACGCTTACCC-3’ and reverse 5’-GCAATAAACAATTAGGTGCTATCCAGGCCCAG-3’).

If not specified, males were used for all experiments in the main text. Phenotyping was performed on both male and female mice.

### Biochemical analysis

Brains and spinal cords were homogenized in lysis buffer [RIPA buffer (Pierce), 2% sodium dodecyl buffer (SDS), protease (Roche) and phosphatase (Thermo Fisher Scientific) inhibitors]. Lysates were sonicated and microcentrifuged for 20 min at 13000 xg at room temperature and soluble fractions collected. Proteins were separated on NuPAGE™ 4 to 12% bis-tris gels (Invitrogen) and transferred to nitrocellulose membranes (Bio-Rad Laboratories). Membranes were blocked in 5% milk in PBS-T (PBS, 0.1% Tween-20) for 1 hour at room temperature. The membranes were incubated overnight at 4 °C with the following primary antibodies: C9orf72 (12E7, kindly donated by Prof. Dr. Manuela Neumann; 1:4 dilution), COL6 (ab182744, Abcam; 1:1000), Calnexin (sc-6465, Santa Cruz Biotechnology; 1:1000 dilution), GAPDH (14C10, #2118, Cell Signaling). After 3 washes in PBS-T, membranes were incubated with secondary HRP-conjugated antibodies for 1 hour at room temperature. After 3 washes in PBS-T, signals were visualized by chemiluminescence (Amersham imager 680) and quantifications performed using ImageJ software.

Meso Scale Discovery (MSD) immunoassays were performed as previously described [63]– [65], using unlabelled anti-poly(GA) antibody (Sigma Aldrich, #MABN889) as capture, and biotinylated anti-poly(GA) (GA5F2, kindly gifted by Prof Dieter Edbauer) as detector.

### Quantitative reverse transcription PCR

Tissues were dissected and flash frozen. Total RNA was extracted with miRNeasy Micro Kit (Qiagen) and reverse-transcribed using SuperScript IV Reverse Transcriptase (Invitrogen) with random hexamers and Oligo(dT)_20_ primers. Gene expression was determined by quantitative real-time PCR using a LightCycler^®^ and SYBR green (Roche). Relative gene expression was determined using the ΔΔCT method. Primers for mouse *C9orf72* are: 5’-TGAGCTTCTACCTCCCACTT-3’ and 5’-CTCTGTGCCTTCCAAGACAAT-3’. Primers to amplify the knock-in sequence are: 5’-GCGGCGAGTGGCTATTG-3’ (primer located within mouse *C9orf72* gene at exon boundary 1-2) and 5’-GGGTAAGCGTAATCTGGAACATC-3’ (sequence within the HA-tag sequence). Primers for mouse *GAPDH* are: 5’-TAGACAAAATGGTGAAGGT-3’ and 5’-AGTTGAGGTCAATGAAGG-3’.

### Immunohistochemistry

Mice were perfused with prechilled phosphate buffered saline (PBS) and then 4% paraformaldehyde (PFA). Spinal cords were dissected and postfixed in 4% PFA at 4°C for 2 hours. After fixation, spinal cords were washed with PBS, allowed to sink in 30% sucrose solution at 4°C, then stored in 0.02% sodium azide at 4°C until further processing. Spinal cords were embedded in optimal cutting temperature (OCT) compound (Tissue Tek, Sakura, Torrance, CA), and 10 μm sections cut with a cryostat (CM1860 UV, Leica Microsystem).

For immunofluorescence, cryosections were washed 3 times in PBS and blocked in 5% BSA, 1% normal goat serum, 0.2% Triton-X in PBS for 1 hour at room temperature. Sections were then incubated with primary antibodies in blocking solution overnight at 4°C. After 3 washes with PBS, sections were incubated for 1 hour at room temperature in blocking solution with secondary antibodies conjugated with Alexa 488, 546, 594, and 633 (Invitrogen). After 3 washes in PBS, sections were mounted with ProLong™ Gold Antifade Mountant with DAPI (Invitrogen).

The primary antibodies used were: IBA1 (019-19741, FUJIFILM Wako Pure Chemical Corporation; 1:500 dilution), GFAP (AB5804, Abcam; 1:500 dilution), CD68 (MCA1957, Bio-Rad Antibodies; 1:200 dilution), COL6 (ab182744, Abcam; 1:200).

Images were taken using a Zeiss LSM 880 confocal microscope or ZEISS Axio Scan.Z1 slide scanner. Image analyses were performed using ImageJ, Imaris software.

### Locomotor, grip strength and body weight assessment

Behavioural tests were undertaken as previously described [59]. Briefly, tests were performed monthly from 3-to 9-months of age, and in 12-18-month-old mice. Motor coordination was measured by rotarod analysis (Ugo Basile). A power calculation using GPower predicted that for an effect size of 10% deviation from the group mean, with a power of 0.85 and an alpha of 0.05, groups sizes of 28 were needed, so we used 14 females and 14 males per group. Mice were trained the week before starting the test. Then, mice received a session which included three trials of accelerated rotarod for a maximum of 300 seconds. Trials started at 4 rpm speed and accelerated up to 40 rpm in 4 min, the final min of the test was performed at 40 rpm. The average of the three trials was used. A grip strength meter (Bioseb) was used to measure forelimb and hindlimb grip strength. The highest muscle force score of three independent trials was used. Body weight was measured weekly from 3-months of age. Mice were randomized into different experimental groups and the operator was blind to genotype.

### Motor neuron counts

10 μm-thick OCT-embedded spinal cord sections were stained with Cresyl Violet and motor neurons located within the sciatic motor pool were counted in each ventral horn, collected every 60 μm of tissue, covering L3 to L5 levels of the spinal cord.

### *In vivo* isometric muscle tension physiology

Isometric muscle tension physiology was performed as previously described [66], [67]. Briefly, under deep anaesthesia (isoflurane inhalation via nose cone), hindlimbs were immobilized and the distal tendons of the TA and EDL muscles of both hindlimbs were exposed and consecutively attached to force transducers in parallel. Sciatic nerves were exposed bilaterally, at mid-thigh level, severed and the distal stumps placed in contact with stimulating electrodes. EDL muscle motor unit number estimates (MUNE) were determined by gradually increasing the amplitude of repeated square wave stimuli, thereby inducing stochastic changes in contractile force. The total number of motor units recruited over the full range of amplitudes was counted for individual muscles and averaged for each genotype.

### Proteomic analysis

Mouse lumbar spinal cords and cortices were solubilized in SDS-Lysis buffer (2% (weight/vol) SDS in 100 mM TEABC supplemented with Roche protease mini and Phos-STOP cocktail tablets). Automated homogenization was performed using the Precellys evolution homogenizer and Bioruptor assisted sonication. Protein estimation by BCA assay and protein quality confirmed by SDS-PAGE. 200 µg of lysate was aliquoted and processed using S-Trap assisted On-column tryptic digestion as described previously (dx.doi.org/10.17504/protocols.io.bs3tngnn). Peptide eluates were then subjected to TMT labelling and High-pH fractionation for LC-MS/MS, a total of 96 fractions were collected and pooled to 48 fractions, vacuum dried and stored at-80°C until LC-MS/MS analysis.

LC-MS/MS analysis: A total of 48 bRPLC fractions were prepared for mass spectrometry analysis using an Orbitrap Tribrid Lumos mass spectrometer in-line with an Ultimate 3000 RSLC nano-liquid chromatography system. The mass spectrometer was operated in a data-dependent SPS-MS3 mode in a top speed for 2 seconds. Full Scan was acquired at 120K resolution at 200 m/z measured using Orbitrap mass analyser in the scan range of 350-1500 m/z. The data dependent MS2 scans were isolated using quadrupole mass filter with 0.7 Th isolation width and fragmented using normalised 32% HCD and detected using an Ion trap mass analyser which was operated in a rapid mode.

Data analysis for mouse tissue: spinal cord raw MS data was searched with Fragpipe software suite (version: 19.1) [68] against the Uniprot Mouse database appended with (GA)400 sequences for C9orf72 and a common contaminant list exists within Fragpipe. FDR was set at 1% for both protein and PSM level. The protein group output files were further processed using Perseus software suite (version: 1.6.15.0) [70] for downstream statistical analysis. Two-sided Welch’s t-Test with 5% FDR multiple-correction was performed to identify differentially regulated proteins. Gene-Ontology analysis was performed on differentially regulated proteins using enrichR software [71].

Gene Ontology analysis was performed on differentially regulated proteins using the software R and the package enrichR [71], [72] (https://ggplot2.tidyverse.org). We assessed the co-occurrence of significant genes from our proteomics analysis with those found to be co-immunoprecipitated with polyGA in two different works [9], [58], based on Drosophila melanogaster and Homo sapiens lines. First, we found mouse orthologues using DIOPT [73], [74], then we represented the co-occurring genes using the software R and the package ggplot2 [72] (https://ggplot2.tidyverse.org).

## Statistical analysis

All data are presented as mean ± standard error of the mean (SEM). Statistical differences of continuous data from two experimental groups were calculated using unpaired two-sample Student’s t-test. Comparisons of data from more than two groups were performed using a one-way-ANOVA followed by Bonferroni correction for multiple comparisons. When two independent variables were available, comparisons of data from more than two groups were performed using a two-way-ANOVA followed by Bonferroni correction for multiple comparisons. Data distribution was tested for normality using Kolmogorov-Smirnov test; when normality could not be tested, we assumed data distribution to be normal. Statistical significance threshold was set at P < 0.05, unless otherwise indicated. Statistical methods were used to predetermine sample sizes.

## Data collection

Data collection and analysis were performed blind to the conditions of the experiments.

## Data Availability

The datasets generated and analysed during the present study are available from the corresponding authors upon request.

Proteomics data from cortex and spinal cord, MS raw data and search output files have been deposited to the PRIDE ProteomeXchange Consortium via the PRIDE partner repository with the dataset identifier PXD047502.

